# Visual search is constrained by the variability of object-category templates

**DOI:** 10.64898/2026.02.02.702780

**Authors:** Susan Ajith, Daniel Kaiser, Lu-Chun Yeh

## Abstract

Real-world visual search is often performed at the category level: we search for shoes or bags without knowing their exact features in advance. This requires categorical search templates that accommodate the inherent variability within the target category. Here, we examine how the variability in search templates across categories constrains visual search performance. We quantify template variability by measuring variability in object drawings from a large online dataset (Experiment 1) and from a controlled lab-based drawing task (Experiment 2) and in turn relate this variability to performance in categorical search. Across both experiments, higher category variability, and thus broader search templates, were associated with slower responses. Moreover, the observer’s most prioritized object template predicted their search performance better than other observers’ templates, indicating that individual differences in template variability shape visual search. Together, our findings demonstrate that naturalistic visual search is governed by structured variability across both object categories and observers.

## Introduction

Whether we are looking for a banana in the kitchen or our keys on a cluttered desk – being able to search efficiently is critical in many everyday situations. Efficient visual search relies on top-down guidance mechanisms that prioritize likely targets for further processing (Wolfe & Horowitz, 2017; Yu et al., 2023). This top-down guidance depends on pre-activating target representations in working memory, known as target templates or attentional templates, which direct our attention towards potential targets (Desimone & Duncan, 1995; Wolfe, 1994).

Given the critical role of target templates in visual search, extensive research has explored what information templates contain and how that information is represented (Geng & Witkowski, 2019; Wolfe, 2021; Wolfe & Horowitz, 2004). Initially, it was assumed that the template is a single, static, and veridical representation of the target (Duncan & Humphreys, 1989; Eriksen, 1953; Green & Anderson, 1956). However, in many everyday situations, we do not exactly know what the target looks like (e.g., when helping a friend find her handbag at a restaurant). More recent models thus propose that templates are flexible and adapt to the information most useful for finding the target in a given situation (Wolfe, 2021; Yu et al., 2022). In situations where detailed visual feature information about the target is unavailable, target templates can be formed on the category level (Nako et al., 2015; Reeder & Peelen, 2013; Yang & Zelinsky, 2009). For instance, when searching for a car, category-level templates contain features like coarse global contours (e.g., a car silhouette) and category-diagnostic object parts (e.g., a wheel) (Reeder & Peelen, 2013).

Yet, the appearance of real-world objects is not equally predictable across categories. Some basic-level categories are defined by a relatively constrained set of feature configurations (e.g., doors, rackets), while others can differ widely in features like shape, color, or texture (e.g., shoes vary considerably between boots, sneakers, or stilettos). This diagnostic variability may determine the precision of the target template across object categories. Object categories with little variability should activate specific and narrow templates that in turn enable efficient search, while categories with greater variability should activate coarse or broad templates that render search less efficient. Importantly, variability in object templates may not only arise from the variability in appearance across a category. Search templates for a category may also vary across individuals, given the unique experience every individual person has garnered across their lifetimes. For instance, when searching for a clock, some observers might predominantly activate representations of an analogue clock (with a round clockface), while others may activate representations of a digital clock (with a rectangular clockface). Particularly for more variably object categories, different appearances may thus be prioritized across observers.

Previous studies examining how template variability influences visual search (Geng & Witkowski, 2019; Witkowski & Geng, 2022; Yu et al., 2022) have typically relied on simple and artificial visual stimuli, where a small number of feature dimensions can be easily manipulated. For example, Geng & Witkowski (2019) manipulated the variance of feature dimensions (color and motion) between the cue and target, and showed that search performance is better predicted with increasing cue-to-target similarity for the low-variance feature dimension compared to the high-variance feature dimension. While these results demonstrate that inherent template variability affects search, they are not readily transferable to real-world objects, which vary along multiple feature dimensions simultaneously and to different extents across categories. Although evidence suggests that if a cue prompts a certain appearance of an object, searching for exemplars with a different appearance is less efficient (Hout & Goldinger, 2015), we do not know what happens to real-world objects that differ along several interacting dimensions. Therefore, it may not only depend on the cue, but also primarily on the object-category.

How, then, can we measure the variability in target templates for real-world object categories? In the current study, we use drawings as a tool to directly probe object-category representations. Drawings allow participants to express object-specific variability in an unconstrained manner, simultaneously conveying multiple relevant feature dimensions. Line drawings preserve the core representational structure of object categories (Fan et al., 2018) and evoke characteristic, category-specific activation patterns in high-level visual cortex, similar to natural object images (Walther et al., 2011; Lowe et al., 2018; Singer et al., 2023). Recent research has employed drawing tasks to visualize individuals’ internal representations, offering insight into how conceptual and perceptual information is represented in the mind (Wang et al., 2024; Bainbridge et al., 2025; Engeser & Ajith et al., 2025). Crucially, drawings also reveal systematic variations across observers: individual drawers emphasize different features of the same object category, reflecting both shared structure and individual differences in an object’s feature distribution. Consistent with this, neural representations are selectively enhanced for scenes that align with participant’s own drawings (Wang et al., 2025). Together, drawings offer a powerful means for capturing individual-specific feature distributions within target templates.

In our experiments, we leverage variability in drawings of real-world object categories as a proxy for variability in the corresponding search templates. The key idea is that categories that yield greater variability in exemplar drawings also afford broader and less precise search templates, whereas categories that yield comparably little variance in exemplar drawings afford narrow and specific search templates.

After quantifying the variability in drawings, we directly tested this idea in two visual search experiments, where participants were cued to search for categorical object targets. Critically, we hypothesized that categories with greater drawing variability and, hence, broader search templates should lead to less efficient search. In Experiment 1, we tested this hypothesis at scale, quantifying variability in search templates for 200 real-world object categories and measuring search performance for these categories. In Experiment 2, we provide a more targeted investigation where we quantify variability for a smaller set of 12 object categories at an individual level and relate object-specific variability in each participant’s own object representations to their search performance.

## Results

### Experiment 1 – Investigating variability in search templates for object categories at scale

In Experiment 1, we examined the effects of template variability on search performance for 200 real-world object categories. Firstly, to quantify template variability, we randomly selected 10 drawings per object category from the THINGS DRAWINGS database (Mukherjee et al., 2025). Ten drawings were chosen because it was the minimum number of available drawings across objects. We computed pairwise similarities between all drawings using a VGG16 DNN trained on object categorization (Deng et al., 2009; Simonyan & Zisserman, 2015; Figure 1A). We extracted activations from an early, intermediate, and late layer along the network (convolutional layers 1_2 and 4_2, and fully-connected layer fc7), yielding a quantification of the drawings’ diagnostic low-level, mid-level, and high-level features (Cichy & Kaiser, 2019). For each layer, we computed pairwise cosine similarity between the ten drawings of each object category. We averaged the resulting pairwise similarities and subtracted them from 1 to yield a measure of variability for each object.

**Fig 1.**
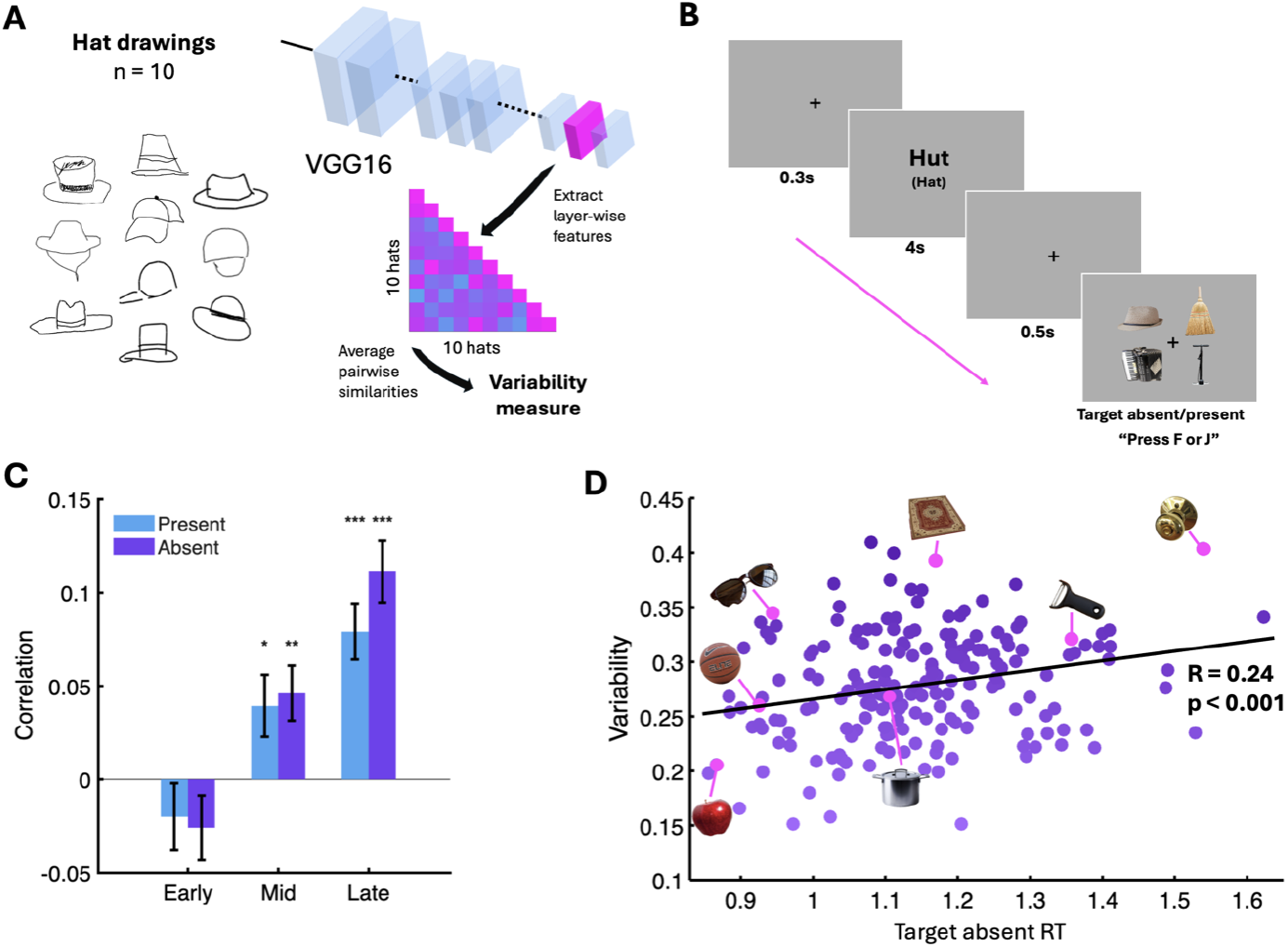
Investigating template variability at scale. (A) Quantification of variability for object categories. First, we selected 10 drawings for each of 200 object categories from THINGS DRAWINGS (Mukherjee et al., 2025). Second, we computed pairwise cosine similarity between these objects using layer-wise activation patterns extracted from a VGG16 DNN. Finally, we averaged the resulting pairwise similarities and subtracted them from 1 to yield a measure of variability for each object category. **(B) Visual search paradigm**. Participants were given a word cue (in German) and asked to report whether the target was present or absent in the following search display. Each object was cued only once per participant. The cue in English was not shown during the experiment. **(C) Correlation between object variability and RT**. For each participant, we correlated the variability score for each object category and their RT in the trial where the category was cued. Variability computed from mid and late layers of the DNN was significantly correlated with search RT in both target-present and target-absent trials, indicating that variability in categorical search templates determines search performance. **(D) Correlation between object variability and average search RT on target absent trials**. For each object, its variability is plotted against its mean RT for target absent trials across participants. Each point represents an object category and representative target images are shown. Error bars indicate SEM. *: p<0.05, **: p<0.01, ***: p<0.001 (FDR-corrected where applicable).

We next conducted a visual search experiment, in which a new set of participants (N=33) performed a cued visual search task involving the same 200 real-world objects (Fig 1B). In each trial, a word cue indicating the target category (in German) was presented for 4s, followed by a search display containing four objects. Participants were instructed to indicate whether the cued object was present or absent. Each target object was cued only once in the experiment. This design, with many objects and no repeats, was chosen to prevent participants from adapting to the object variability statistics of the target images shown during the experiment. We hypothesized that search performance, measured using response times (RTs) should depend on the variability of the categorical target templates – quantified using the variability in exemplar drawings.

#### Variability in search templates for object categories constrains search performance

Our analysis focused on RTs given the high overall task accuracy (M=94.6%, SD=3.2%). We correlated each participant’s RT for the target present and absent trials per object category with the objects’ variability score extracted from the DNN (Fig 1C). Critically, search RT varied significantly with drawing variability computed from DNN activations in the intermediate and late layers. This effect was evident for both target-absent trials (late: *r*=0.11, *t*(32)=6.78, *p*<0.001, intermediate: *r*=0.046, *t*(32)=3.13 *p*<0.001; FDR-corrected) and target-present trials (late: *r*=0.079, *t*(32)=5.28, *p* <0.001, intermediate: *r*=0.039, *t*(32)=2.37, *p*=0.012; FDR-corrected), indicating faster search for object categories with less variable templates, whereas greater template variability led to slower search. No similar effects were found for the early DNN layer (target-absent: r=0.026, t(32)=1.49, p=0.92; target-present: r=0.02, t(32)=1.10, p=0.861), indicating that variability in visual features of intermediate-to-high complexity constrains target templates. To illustrate the relationship between template variability and search RT, we further correlated the mean RT in target-absent trials for each object across participants with the objects’ variability score, again yielding a robust relationship (*r* = 0.241, *p* <0.001) (Fig 1D). Together, our results demonstrate that variability in target templates for real-world object categories shapes search behavior, with less efficient search for categories with more variable mental templates. In Experiment 2, we further scrutinized this effect by quantifying template variability on the individual-participant level and measuring search performance in the same participants.

### Experiment 2 – Investigating variability in search templates for object categories at the individual level

In Experiment 2, we investigated how variability in object templates impact search on the individual level. The goal of this experiment was twofold: First, we aimed to provide a replication of Experiment 1, with a smaller set of objects and with a quantification of variability that rests on drawing variability within participants. Second, we aimed to clarify whether individual differences in object templates are capable of explaining individual differences in search performance. To enable this, we focused on a smaller set of 12 real-world object categories and quantified their variability for each individual participant. Our participants (N=31) performed two tasks: First, they performed a drawing task to capture the variability in mental representations for the 12 object categories, where greater variability among drawings indicates a broader target template. Second, the same participants performed a cued visual search task to quantify how search performance varies as a function of template variability.

During the drawing task, participants were given 1 minute to create four drawings for each of the 12 object categories on an Apple iPad Pro using an Apple Pencil. Each page contained a 2×2 grid with the object name (in German) written in the center and they had to draw the objects in a fixed order (1-4) (See Fig 2A). To quantify the variability of drawings, we followed the same DNN-based procedure as outlined for Experiment 1. Here, given that we only had relatively few object categories and individual drawings, we included a second step: As our VGG16 DNN was pretrained on real-world object images, we replaced participants’ drawings with hand-picked real-world images that closely matched each drawing (Fig 2A). The additional step of hand-picking suitable object photographs indeed yielded better predictions of search performance, compared to the DNN evaluating the drawings directly (see Supplementary Figure S3). Further, we only used the penultimate DNN layer, given that it yielded the most robust results in Experiment 1 (Fig 1C, Supplementary Fig S1).

**Fig 2.**
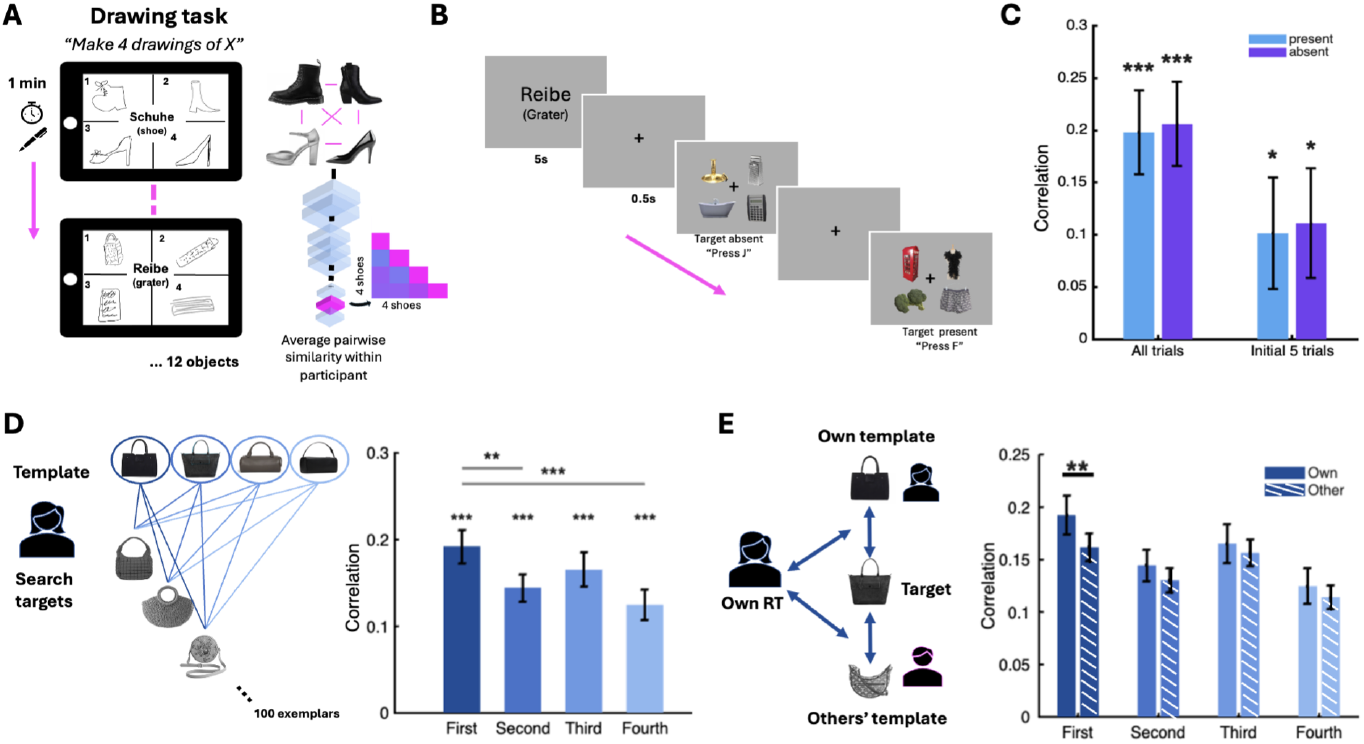
Investigating template variability at the individual level. (A) Quantification of variability for object categories. Participants first produced four exemplar drawings for each of 12 real-world object categories. Here, participants’ drawings were replaced with hand-picked real-world images that closely matched each drawing and variability was then quantified using the same DNN-based pipeline as in Fig. 1A, yielding object-specific variability scores for each object category and each individual participant. **(B) Visual search paradigm**. After the drawing task, the same participants that made the drawings completed a cued visual search task. In each block, participants first saw a word cue and on all subsequent trials responded whether the corresponding target category was present or absent. **(C) Correlation between object variability and RT**. For each participant, we correlated the variability scores and mean search RT per object. Objects with more variable drawings yielded slower responses, in both target-present and target-absent trials. This effect was already significant when restricting analyses to the first five trials for each search target, indicating that template variability influenced search from the outset. **(D) Correlation between template-to-target similarity and RT**. Given that participants produced four drawings per category in a fixed order, we tested whether the drawing order captured the priority of certain object appearances in the search template. Using a VGG16, we computed cosine similarity between each of the objects selected to be similar to the four drawings and all target objects shown during the experiment. The targets’ similarity to all four drawings significantly predicted search RT, with the first drawing showing the strongest correlation, suggesting that different appearances of an object are weighted in the search template according to their likelihood. **(E) Correlation between individual target similarity and RT**. For each participant, we computed the cosine similarity between the VGG16 activation vectors for each participant’s own drawings and activation vectors for all target images of the corresponding category in the search task. We then correlated these similarity values and that participant’s RTs in the corresponding target-present trials (within-participant correlation, “own” condition) and compared it to the same correlations when other participants’ drawings were used (between-participant correlation, “other” condition). We observed significantly greater correlations in the own condition compared to the other condition when the first drawing was used, suggesting that individual target templates induce idiosyncrasies in search performance. Error bars indicate SEM. *: p<0.05, **: p<0.01, ***: p<0.001 (FDR-corrected where applicable).

After the drawing task, participants performed the cued visual search task. Here we used a block design, where the search target remained the same for each block of 100 trials. In the beginning of each block, a word cue (in German) indicated the target for the entire block. Each trial then featured a search display containing four objects. Participants were asked to indicate whether the cued object was present or absent.

#### Replication: Variability in templates constrains search performance

To test how template variability impacts search performance, we again correlated the variability in exemplar drawings and the search performance across object categories. This analysis essentially replicates Experiment 1. Here, however, given that we estimated the variability among exemplar drawings stemming from individual participants, we quantified template variability separately for each participant and predicted their own search data from this variability.

Our analysis again focused on search RTs given the high overall task accuracy (M=94.8%, SD=2.2%). Critically, search RT varied significantly with template variability, both on target-absent trials (*r* =0.206, *t*(30) = 5.11, *p*<.001; FDR-corrected) and on target-present trials (*r*=0.198, *t*(30)=4.93, *p*<.001; FDR-corrected). Despite the high overall accuracy, we also observed a similar overall pattern when correlating error rates with template variability, which was significant for the target absent trials (r=0.115, t(30)=3.71, p<0.001) but only numerically for the target-present trials (r=0.047, t(30)=0.924, p=0.181) (see Supplementary Fig S4). These results indicate that narrower search templates speed up search, whereas broader templates lead to slower search, replicating the pattern from Experiment 1.

Additionally, we tested whether template variability impacted search from the outset, before participants could fully estimate the variability of actual target images in the experiment. To this end, we analyzed only the first five target-present and target-absent trials for each object category. Crucially, the correlation between template variability and search RT was significant even when only these initial trials were considered, for both target-absent trials (*r*=0.11, *t*(30)=2.11, *p*=0.033) and target-present trials (*r*=0.10, *t*(30)=1.90, *p*=0.040). This suggests that real-world object variability impacted search from the outset – much like in Experiment 1, where each object was only cued once and no learning of target distributions was possible.

Next, we take a closer look at how the similarity between individual drawings made by the participants and the search targets presented on every trial shapes search performance. These analyses provide insights into how search templates are prioritized at the individual level.

#### Object appearances are prioritized differently in the search template

We investigated how participants prioritised different possible appearances of an object category in their search template. Given that participants produced 4 drawings per object in a fixed order, we reasoned that this order might reflect the relative priority of different object appearances. For example, drawings of a clock varied across participants: some drew an analogue clock first and a digital clock later, whereas others did the opposite. This implies that the highest-priority appearance for a “clock” could be an analogue clock for some individuals but a digital clock for others. Based on this reasoning, we hypothesized that objects drawn earlier should yield higher priorities, yielding faster search times when the target on a given trial more closely matches the object that was drawn first. For instance, a participant who prioritises an analogue clock should show the fastest search on trials where the target resembles an analogue clock (e.g., a round clockface). To quantify template-to-target similarity, we extracted DNN activation vectors for all template images (i.e., objects selected to be similar to the four drawings) and all target images (i.e., target objects shown during the experiment) from the penultimate layer of the VGG16. We then computed the cosine similarity between the activation vectors for each target object and the four template images. Finally, for each participant, we correlated the template-target similarity and the participant’s RT in the corresponding target-present trials. This was done separately for each of the four exemplar drawings, yielding a measure of how similarity to each possible object template impacts search performance (Fig 2D).

Distance to the four drawings significantly correlated with search RTs, with the first drawing showing the strongest correlation (first: *r*=0.21, *t*(30)=12.65, *p*<0.001; second: *r*=0.16, *t*(30)=11.27, *p*<0.001; third: *r*=0.18, *t*(30)=10.14, *p*<0.001; fourth: *r*=0.14, *t*(30)=7.97, *p*<0.001; FDR-corrected). Correlations were significantly stronger for the first than for the second and fourth drawing (first vs. second: diff=0.053, t(30)=4.48, p<0.01; first vs. fourth: diff=0.073, t(30)=5.12, p<0.001; FDR-corrected). The difference between the first and third template did not reach significance (diff=0.037, t(30)=2.28, p=0.06). Together, these results suggest that object appearances that are more salient, and thus drawn first, are prioritized in the search template.

#### Individual differences in target templates explain individual differences in search

Finally, we tested whether participants’ own drawings preferentially predict their own search performance. We reasoned that if this was the case, the similarity of search targets to their own object exemplar drawings should predict their search performance better than the similarity of the search targets to other participants’ drawings. Returning to the analogue–digital clock example, a participant who prioritises analogue clocks should search fastest when the target resembles their own analogue template (e.g., a round clockface), rather than someone else’s digital template (e.g., a rectangular display). Evidence for this pattern would indicate that participants rely on their *own* object templates to guide attention. Because similarity to the first drawing was the strongest predictor of performance at the group level, we expected these individual differences to be most pronounced when using distances to the first drawing to predict search performance.

To test this, we computed the cosine similarity between the VGG16 activation vectors for participants’ *own* template and activation vectors of all target images of the corresponding object in the search task. We then correlated the template-target similarity values and the participant’s RTs in the corresponding target-present trials, yielding a measure of the search performance relative to each participant’s *own* template. Next, to assess whether the effect on search performance is uniquely influenced by the similarity of the search target to participants’ own exemplar drawings, we repeated this analysis using *other* participants’ exemplar drawings. For each participant, we calculated the cosine similarity between activation vectors of *others’* templates and target images. Then, we correlated *others’* template-target similarities and the participants’ *own* RTs in the corresponding target-present trials. The resulting similarity–RT correlations were averaged across participants. This analysis was repeated for all four exemplar drawings for each of the 12 objects. We thus obtained two measures for each object template: 1. Within-participant correlations, capturing how search behavior is predicted by the similarity of each target and participants’ own drawings, and 2. between-participants correlations, capturing how search behavior is predicted by the similarity of each target and other participants’ templates. Comparing the within- and between-participant correlations, we quantified how much an individual participant’s search performance is shaped by how similar the target is to their own personal search templates.

The similarity of the search targets to participants’ own first drawing of each object correlated significantly more with their search performance than similarity to other participants’ first drawings (diff=0.034, t(30)=3.02, p<0.01). Although similar numerical trends were observed for the second to fourth drawings, the differences did not reach significance (second: diff=0.016, t(30)=1.13, p=0.26; third: diff=0.010, t(30)=0.62, p=0.27; fourth: diff=0.011, t(30)=0.76, p=0.27; FDR-corrected). These results indicate that targets that are more similar to a participant’s own first drawing, and thus their own most salient appearance of an object, were associated with faster search performance. Individual differences in search performance are thus partly shaped by individual differences in the most prioritized object appearance in each participant’s personal search template.

## Discussion

Object categories differ substantially in their visual appearance and each category is defined by its own distribution of features (Ullman, 1989). To successfully search for objects in the world, individuals must therefore represent the inherent variability in feature distributions to adapt to different instances of an object. Our assumption is that objects that have greater variability amongst exemplars afford broader and less precise search templates, whereas categories that yield comparably little variance amongst exemplars afford narrow and specific search templates. By evaluating similarities among multiple exemplar drawings of an object category, we quantified the variability in mental templates, and then relate this variability to visual search performance. Across two experiments, we show that search is less efficient for categories with more variable mental representations and more efficient for categories with less variable representations. In Experiment 1, we quantified variability at scale across 200 real-world object categories from the THINGS DRAWINGS database (Mukherjee et al., 2025) and independently measured search performance. We demonstrate that inherent object variability systematically correlates with search performance across object categories. In Experiment 2, we quantified object variability and probed search performance in the same individuals. We replicate the results from Experiment 1 and additionally show that individual differences in prioritizing likely object appearances predict individual differences in search performance. These individual differences in search performance were driven by the most prioritized object appearance within the template (i.e., participants’ first drawings). Together, these findings provide robust evidence that search templates are shaped by the variability of the target category template as well as by the variability of this template across participants.

Real-world visual search is often considered surprisingly efficient when compared to search in simplified displays (Peelen & Kastner, 2014; Wolfe et al., 2011). However, much of the literature has focused on object categories with relatively narrow and stable templates, such as human bodies, faces, and cars (Peelen et al., 2009; Reeder & Peelen, 2013; Simpson et al., 2014). Yet, real-world search may proceed differently depending on what is being searched for. While searching for people or cars is easy due to their relatively consistent visual appearances, searching for more variable objects such as shoes or bags, can be a lot harder. Greater variability in object-category appearances likely produces less precise and more uncertain target templates, which in turn can influence both attentional guidance and target processing (Kristjánsson, 2023; Witkowski & Geng, 2023). Future work could examine how inherent object variability systematically shapes visual search by explicitly treating object-category variability as a fundamental factor in visual search designs.

In the present study, we probe search templates grounded in long-term visual experience, rather than explicitly manipulating target statistics within the task, as done in prior work (e.g., Carlisle et al., 2011; Geng & Witkowski, 2019; Goldstein & Beck, 2018; Grubert et al., 2016; Yu et al., 2022). Our results therefore indicate that real-world search is also influenced by long-term visual experience rather than by task-specific target statistics. In Experiment 1, this is evident from our one-trial-per-object design, which required participants to update their templates continuously and prevented them from learning within-category variability on the go. In Experiment 2, the critical effect replicated when analyzing only the first few trials of the object category, before participants could effectively learn the distribution of exemplars within a category. In real-world search, such long-term priors are probably balanced against current target distributions in each context. More research is needed to understand how search templates are updated when target statistics change in the real-world, and how such updating affects the balance of long-term experience versus short-term adaptive learning in guiding attention.

Our study also touches upon two critical open questions in the visual search literature. First, we find differences in search performance as a function of object-category variability. This leaves open the question of whether object variability affects template precision during the selection or the identification stage of the search process (Yu, Hanks, et al., 2022). In Guided Search 6.0, Wolfe (2021) distinguishes between a guiding template used during the selection stage, and a target template used during the identification stage, proposing that guidance and identity decisions rely on different informational content and levels of template precision to satisfy distinct computational goals. Although our results do not allow us to directly dissociate these stages, object-category variability may plausibly influence both (Geng & Witkowski, 2019). In particular, because the upper limit of guidance is constrained by the precision of the target template stored in memory (Yu et al., 2023), broader object-category templates may lead to less efficient selection via the selection of multiple potential target candidates, prolonged identification to eliminate distractors, or a combination of the two. Second, we observe that participants prioritize different appearances of the same object category, raising the question of whether search relies on multiple templates or on different instantiations of a single, flexible object template (Yu et al., 2023; Yu, Johal, et al., 2022). While this cannot be resolved directly with our data, our findings suggest that in real-world visual search, the distinction between multiple versus single templates may be especially relevant for highly variable object categories. One promising paradigm to address this question is visual foraging (Wolfe, 2013; Wolfe et al., 2016; Hong & Wolfe, 2025), which allows investigation of search behavior when multiple instances of a target category are simultaneously present. In this setup, one could examine whether observers rely on multiple templates, leading to sequential harvesting of similar-looking exemplars before moving onto the next object appearance, or on a single broad template, resulting in more variable harvesting patterns across object appearances.

We find that object-category templates differ systematically across individuals. Previous work has shown that search preparation involves the priming or preactivation of neuronal populations that are selective to the target category (Chelazzi et al., 1993, 1998; Desimone, 1998; Peelen & Kastner, 2011, 2014), and this is most effective when it is directed toward the most distinguishable target representation (Battistoni et al., 2017). However, what constitutes the most distinguishable object representation can differ across individuals based on variations in their visual experience or “visual diet” (Barrett, 2020; Hartley, 2022), as well as from individual differences in visual system architecture (Kanai & Rees, 2011; Moutsiana et al., 2016). Consistent with this view, our results reveal individual differences in search performance, where people use their own personal object-category templates to guide attention. Search was faster when the target more closely matched an individual’s most prioritized object appearance (i.e., their first drawing) compared to other participants’ most prioritized appearance (i.e., someone else’s first drawing). This shows that not all individuals converge on the same single prototypical exemplar for an object category. Particularly when it comes to the most likely appearance of the targets, individual differences in this most prioritized appearance drive differences in target detection during search.

While individual differences were strongest for the first drawing, corresponding to the most prioritized object appearance, a numerical trend was also observed for later drawings. How such effects play out may depend on the object category. When analyzing individual differences at an object-level (see Supplementary Fig. S5), we made two interesting observations: First, some categories showed weaker individual differences for the first drawing but stronger individual differences for subsequent drawings. For example, in the case of ice cream, the most prioritized template may correspond to a highly shared prototype (e.g., an ice cream cone), whereas variability might emerge in later drawings which depicts less typical but more personally salient exemplars. Other objects in our set may not have a well-agreed on most typical example, so that the effect was most pronounced for the first drawings overall. Second, we found a positive correlation between category variability (i.e., the variability estimated by the DNN; Fig. 2A) and the magnitude of individual differences (i.e., the mean differences between the own and other conditions across all drawings per category; Fig. S5) across the 12 categories (r=0.65, p=0.02). This suggests that object categories with higher exemplar variability yield more individualized search templates.

Our findings point to three promising avenues for future research. First, prior work in object recognition has shown substantial individual differences in general recognition ability (Gauthier & Tarr, 2016; Richler et al., 2019). These differences have been linked to variations in mental prototypes and representational structure, raising the possibility that individual differences in search templates may relate to differences in core object-recognition abilities. An important open question, therefore, is whether the personalized search templates observed here generalize across tasks and predict individual performance in object recognition more broadly. Second, variability constraints across object categories and observers may also influence performance in other capacity-limited tasks like visual short-term memory (Posner & Keele, 1968; Bays et al., 2024) and determine that the magnitude of neural competition in visual cortex (Cohen et al., 2016; Cohen, 2019; Kaiser et al., 2019). Finally, our results demonstrate that even a limited number of participant-generated drawings can be used to predict search performance, highlighting drawings as a flexible and efficient tool for probing mental representations and their variability (Fan et al., 2023; Bainbridge et al., 2025; Engeser & Ajith et al., 2025). Future work could further explore the potential of drawings to predict performance across object categories, tasks, and observers in visual search and beyond.

Taken together, our findings show that real-world visual search is constrained by the inherent variability of object-category templates as well as by individually specific variability constraints arising from prior world knowledge. Search is therefore neither uniform across objects nor across observers, and individual experience with object variability influences how quickly an object can be found. This realization calls for theoretical frameworks that appropriately honour the idiosyncratic nature of mental templates for real-world objects. Such frameworks promise a leap forward in how well we can predict, and explain, how we search in everyday situations.

## Methods

### Experiment 1

#### Participants

Thirty-four healthy native German speakers (16 females, mean age=24.7y, SD=5.1y) participated in the experiment. Sample size was determined to achieve ∼80% power for detecting a hypothetical medium-sized effect of d=0.5 at p<.05 (two-tailed t-test). The participants provided informed written consent and received a compensation of 10 euro per hour for their time. The study was approved by the Ethics Committee of the Julius Liebig University Giessen and was in accordance with the 6th Declaration of Helsinki. One participant data was invalid because the file was not saved correctly.

#### Stimuli

Two-hundred inanimate objects were selected as targets from the THINGS image database (Mukherjee et al., 2025). Another 500 objects were selected as distractors. Familiarity ratings for all objects were acquired from THINGS Drawings recognition data (Mukherjee et al., 2025). Only objects which were familiar to more than 90% of raters were included as targets. All stimuli were fitted onto a transparent square background.

#### General Experimental Procedure

Participants were first presented with a list of the 200 target object names in German, to ensure that they were familiar with all object names. Participants then received written and verbal instruction about the cued visual search task, followed by 10 practice trials using target objects that did not appear in the main experiment. Participants were instructed to respond as quickly and accurately as possible.

#### Visual search task

The cued visual search task was programmed in PsychoPy v2024.2.2 (Peirce et al., 2019). On each trial, a word cue indicating the target category (in German) was presented for 4s. Each target object was cued only once in the experiment, resulting in 100 target-present and 100 target-absent trials (200 trials total). This design, with many objects and no repeats, was chosen to prevent participants from learning object variability statistics from the targets. After a central fixation cross presented for 1s, a search display containing four objects was presented. Each object subtended approximately 3×3 degrees visual angle and was presented at approximately 2.8 degrees eccentricity. Participants were instructed to indicate via a keyboard button whether the cued object was present (“F”) or absent (“J”). Assignment of target-present and target-absent trials was counterbalanced across participants, as were target locations and keyboard responses. Trial order was randomized across participants. The trials were divided into two blocks, with a self-paced break in between.

After completing the search task, participants filled out a questionnaire rating all target objects on two aspects: 1. Variability, indicating how much an object can vary in appearance (1–10 scale), and 2. Familiarity, indicating how often they encounter the object in daily life (1–10 scale). The total duration of the experiment, including the search task and questionnaires, was approximately 60 minutes per participant.

#### Data analysis

We assumed that the variability of object drawings across individuals captures the real-world variability of an object category. To quantify this variability, first we randomly selected 10 drawings per object category from the THINGS DRAWINGS database (Mukherjee et al., 2025). We employed a deep neural network (VGG16, pretrained on object recognition) (Deng et al., 2009; Simonyan & Zisserman, 2015) to measure the similarity between the ten drawings for each of the 200 objects. For each object drawing, activation vectors were extracted from the convolutional layer 1_2, convolutional layer 4_2, and fully-connected layer fc7 of the network. We focused on these layers because it provides an approximation from low-level to complex feature processing, similar to how drawings tend to carry abstract ions of high-level object features. For each layer, pairwise cosine similarity between the ten drawings of each object was computed. The resulting pairwise similarities were averaged and subtracted from 1 to yield a measure of variability for each object.

Next, we investigated the relationship between template similarity and search performance. Both accuracy and response times (RTs) were examined, with RT analyses restricted to correct trials. For each participant, we computed the Pearson correlation between the variability score and mean RT per object from the search task. These correlations were calculated separately for target-present and target-absent trials, as well as for each layer. Correlations across participants were then tested against zero using one-tailed one-sample t-tests, and false discovery rate (FDR) correction was applied to control for multiple comparisons.

### Experiment 2

#### Participants

Thirty-four healthy native German speakers (24 females, mean age = 24.78, SD = 3.15 years) participated in the experiment. Sample size was determined to achieve ∼80% power for detecting a hypothetical medium-sized effect of d=0.5 at p<.05 (two-tailed t-test). All participants had normal or corrected-to-normal vision. They provided informed written consent and received a compensation of 10 euro per hour for their time. The study was approved by the Ethics Committee of the Julius Liebig University Giessen and was in accordance with the 6th Declaration of Helsinki. Three participants were excluded from the analysis due to noncompliance with the task instructions, as they produced identical drawings for all objects.

#### Stimuli

Twelve everyday objects were selected as targets. For each target, we collected 100 exemplar images selected from Google Images. Another 350 stimuli selected from (Konkle & Oliva, 2011) were used as distractors, with only inanimate and non-vehicle object images used. All stimuli were fitted onto a transparent square background.

#### General Experimental Procedure

The experiment was conducted in two parts - drawing session followed by the search task.

##### Drawing task

Participants were asked to create four drawings for each of twelve object categories (Apple, Basketball, Ice Cream, Handbag, Lamp, Razor, Grater, Shoe, Clock, Racket, Ventilator, and Door) on an Apple iPad Pro using an Apple Pencil via the Sketchbook App. Each page contained a 2×2 grid with the object name (in German) written in the center. Participants had 1 minute per object to complete the four drawings, following the order 1–4. The order of objects was randomized across participants.

##### Visual search task

After the drawing task, participants received written and verbal instructions for the cued visual search task, which was programmed in PsychoPy v2024.2.2 (Peirce et al., 2019). The task consisted of 24 blocks divided into two sessions, with each session containing one block for each target object. The order of blocks within each session was randomized. Each block contained 100 trials (50 target-present, 50 target-absent), resulting in 2,400 trials in total. At the start of each block, a word cue indicating the target for the entire block (in German) was presented for 5s. Each trial began with a central fixation cross for 1s, followed by a search display containing four objects. Participants completed 12 practice trials searching for objects that were not targets in the main experiment to familiarize themselves with the instructions and procedure. During the main task, participants indicated whether the cued object was present (“F”) or absent (“J”) and were instructed to respond as quickly and accurately as possible. Target locations were balanced across trials, and keyboard responses were counterbalanced across participants. Trial order was randomized within blocks. Each target object appeared only once in the experiment. Participants could rest between blocks and initiate the next block at their own pace.

### Data Analysis

#### Relationship between template variability and search performance

To quantify drawing variability, we followed the same DNN-based procedure as outlined for Experiment

1. However, given that we only had relatively few object categories and individual drawings, we included a second step: As our VGG16 DNN was pretrained on real-world object images, we replaced participants’ drawings with hand-picked real-world images that closely matched each drawing. All images were converted to grayscale and resized before passing through the DNN (Willenbockel et al., 2010).

Next, we investigated the relationship between template similarity and search performance. Both accuracy and reaction times (RTs) were examined, with RT analyses only restricted to correct trials. For each participant, we computed Pearson correlation between the variability scores and error rate/mean RT per object from the search task. These correlations were calculated separately for target-present and target-absent trials. Additionally, we analyzed the first-five target-present and target-absent trials, to test whether template variability impacted search from the outset, before participants could estimate the variability of actual target images in the experiment. Correlations across participants were then tested against zero using one-tailed one-sample t-tests, and false discovery rate (FDR) correction was applied to control for multiple comparisons.

#### Priority of multiple templates in search performance

To quantify template-to-target similarity, activation vectors for all template images (i.e., objects selected to be similar to the four drawings) and all target images (i.e., target objects shown during the experiment) were extracted from the penultimate layer of the VGG16 DNN. We then computed the cosine similarity between the activation vectors for each target object and the four template images. Finally, for each participant, we computed Pearson correlation between the template-target similarity values and the participants’ RT in the corresponding target-present trials. Correlations across participants were then tested against zero using one-tailed one-sample t-tests, and false discovery rate (FDR) correction was applied to control for multiple comparisons.

#### Relating individual differences in target templates and visual search

For each participant, we computed the cosine similarity between the VGG16 activation vectors for participants’ *own* template and the activation vectors of all target images of the corresponding object in the search task. We then correlated the template-target similarity values and the participant’s RTs in the corresponding target-present trials, yielding a measure of the search performance relative to each participant’s *own* template. Next, we repeated this analysis using *other* participants’ exemplar drawings. For each participant, we calculated the cosine similarity between activation vectors of *others’* templates and target images. Then, correlated *others’* template-target similarities and the participants’ *own* RTs in the corresponding target-present trials. The resulting similarity–RT correlations were averaged across participants. This analysis was repeated for all four exemplar drawings for each of the 12 objects. We thus obtained two measures for each object template: 1. Within-participant correlations, capturing how search behavior is correlated with participants’ *own* template, and 2. between-participants correlations, capturing how search behavior is correlated with *others’* templates. By subtracting the between-participant correlations from the within-participant correlations, we quantified how much an individual participant’s own search template is uniquely associated with their search behavior across object exemplars. The difference between within and between participants’ correlations were tested against zero using one-tailed one-sample t-tests, and FDR correction was applied to control for multiple comparisons.

## Author contributions

Conceptualization: S.A., D.K., and L.-C.Y. Methodology: S.A. and L.-C.Y. Investigation and formal analysis: S.A. Visualization and writing-original draft: S.A. Writing-review and editing: S.A., D.K., and L.-C.Y. Supervision: D.K. and L.-C.Y. Funding acquisition: D.K. and L.-C.Y.

## Funding

S.A. is funded by a Graduate Scholarship from the Justus Liebig University Giessen. D.K. is supported by the DFG (KA4683/5-1, project number 518483074; KA4683/6-1, project number 536053998; KA4683/7-1, project number 548389777) and an ERC Starting Grant (PEP, ERC-2022-STG 101076057). L.-C.Y. is supported by the MSCA programme (101149060). This work is further supported by the DFG under Germany’s Excellence Strategy (EXC 3066/1 “The Adaptive Mind”, project number 533717223). Views and opinions expressed are those of the authors only and do not necessarily reflect those of the funders. Neither the funders nor the granting authority can be held responsible for them.

We thank Malaika Grace Alphonsus and Sirine Nouira for assistance with data collection and image selection.

## Data availability

Data and code are available through the project’s OSF page (https://osf.io/e2qk8).

## Supplementary Material

**Table S1:** Experiment object list. Words were presented in German in the experiment.

|  |  |  |  |  |  |  |  |
| --- | --- | --- | --- | --- | --- | --- | --- |
| Pan | Car | Jellyfish | Paper plate | Plant | Anchor | Kettle | Comb |
| Bag | Couch | Drum | Watch | Doorknob | Motorcycle | Guitar | Flashlight |
| Airplane | Chess piece | Snowman | Coffee pot | Coffin | Medal | Headphones | Blender |
| Glasses | Notebook | Tree | Croissant | Recliner | Perfume | Helicopter | Sponge |
| Purse | Bottle | Briefcase | Helmet | Jersey | Candle | Toothbrush | Racket |
| Sofa bed | Glass | School bus | Backpack | Bone | Sailboat | Lemon | Cup |
| Measuring cup | Jellyfish | Polo shirt | Tent | Apron | Pacifier | Pedal | Cactus |
| Rug | Drum | Dumbbell | Umbrella | Bouquet | Table | Keyboard | Bowl |
| Locket | Snowman | Rocket | Banana | Swimsuit | Log | Pen | Pomegranate |
| Computer | Tree | Gravestone | Eraser | Dishwasher | Flask | Envelope | Shoe |
| Gun | Briefcase | Hairbrush | Lightbulb | Doormat | Wallet | Syringe | Pencil sharpener |
| Leaf | School bus | Hat | Spoon | Dollhouse | Teapot | Fence | Coconut |
| Water bottle | Polo shirt | Crown | Mixer | Globe | Record player | Sunglasses | Pocket watch |
| Garbage truck | Dumbbell | Screwdriver | Water fountain | Camera | Apple | Gingerbread man | Ice cream |
| Rocking chair | Rocket | Taco | Jet | Pipe | Laundry basket | Cake | Toy |
| Spatula | Gravestone | Skateboard | Feather | Trophy | Pencil | Clock | Bucket |
| Knife | Hairbrush | Donut | Ship | Scooter | Boot | Scarecrow | Ring |
| Dice | Hat | Cage | Radio | Bus | Stethoscope | Piggy bank | Microphone |
| Bed | Crown | Crayon | Frame | Speaker | Fan | Razor | Jug |
| Television | Screwdriver | Bread | Ladder | Cellphone | Palm tree | Pot | Flip flop |
| Sock | Taco | Handcuff | Doorbell | Mop | Hourglass | Hairdryer | Jigsaw puzzle |
| Steering wheel | Skateboard | Cookie | Jewelry | Microwave | Suitcase | Glove | Rocking horse |
| Cassette | Donut | Minivan | Nail clippers | Parachute | Piano | Mannequin | Door |
| Tiara | Cage | Boat | Gate | Treadmill | Lamppost | Nest | Punching bag |
| Crayon | Lamp | T-shirt | Painting | Coffee pot | Basketball | Jeans | Jacket |
| Paper plate | Jeep | Coffee table | Peeler | Croissant | Ambulance | Fire hydrant | Mirror |
| Watch | Grater | Cupcake | Gift | Helmet | Traffic light | Scissors | Sandal |
| Acorn | Bench | Abacus |  |  |  |  |  |

**Table S2:** Experiment 2 object list. Words were presented in German in the experiment.

|  |
| --- |
| Apple |
| Racket |
| Basketball |
| Icecream |
| Razor |
| Grater |
| Door |
| Clock |
| Ventilator |
| Bag |
| Shoes |
| Lamp |

**Fig S1:**
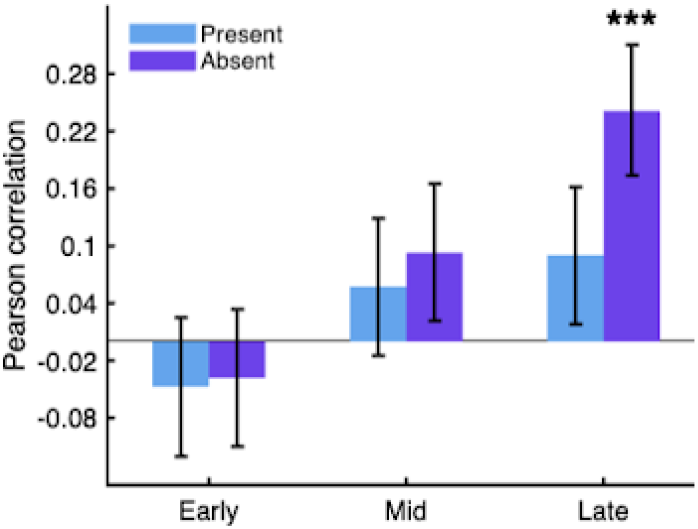
Correlation between object variability and mean RT across participants in Experiment 1. Pearson correlation was calculated between variability score of each object and its mean RT across participants. Search time varied significantly with object variability for target-absent trials for the penultimate layer of VGG16. *: p<0.05, **: p<0.01, ***: p<0.001. (FDR-corrected where applicable).

**Fig S2.**
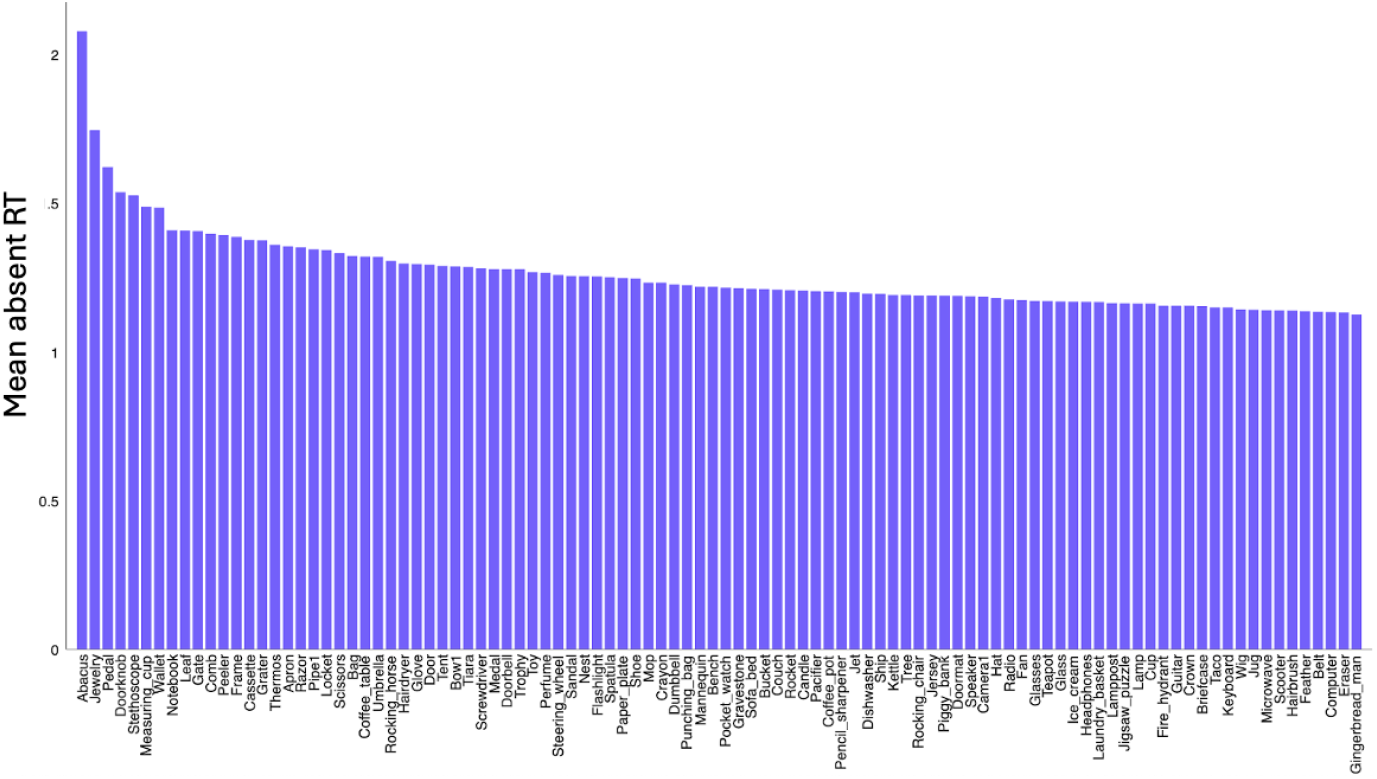
Mean RT for each object category in Experiment 1. Search time of target absent trials for each object was averaged across participants and ranked along the x-axis.

**Figure S3.**
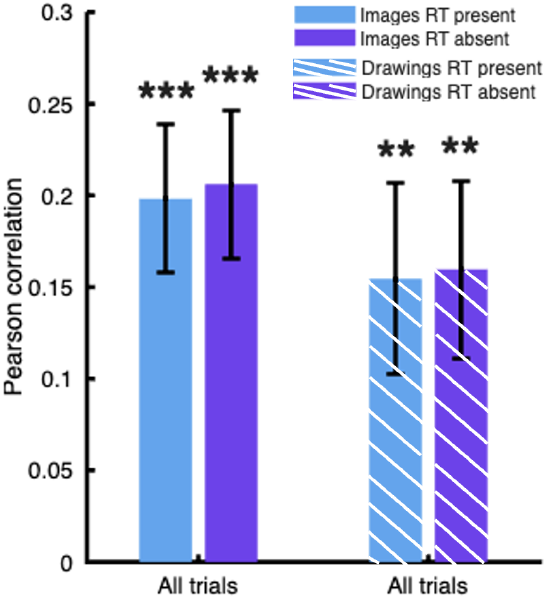
Computing variability from drawings versus images in Experiment 2. Comparison of the correlation between participants’ drawing variability and their search time computed from fc7 DNN activations for original drawings versus the same correlation computed from fc7 DNN activations for real-world photographs similar to these drawings. Error bars indicate SEM. *: p<0.05, **: p<0.01, ***: p<0.001 (FDR-corrected where applicable)

**Fig S4.**
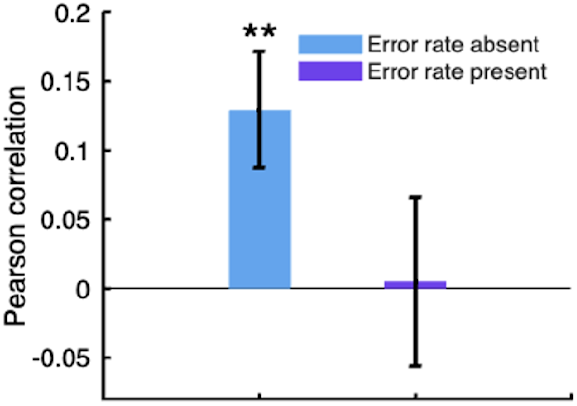
Correlation between participants’ error rate and search time in Experiment 2. Error rate was calculated as the percentage of errors in the search task for each participant. Pearson correlation was calculated between their drawing variability and error rate. Error rate significantly varied with search time for target-absent trials. Error bars indicate SEM. *: p<0.05, **: p<0.01, ***: p<0.001 (FDR-corrected where applicable)

**Fig S5.**
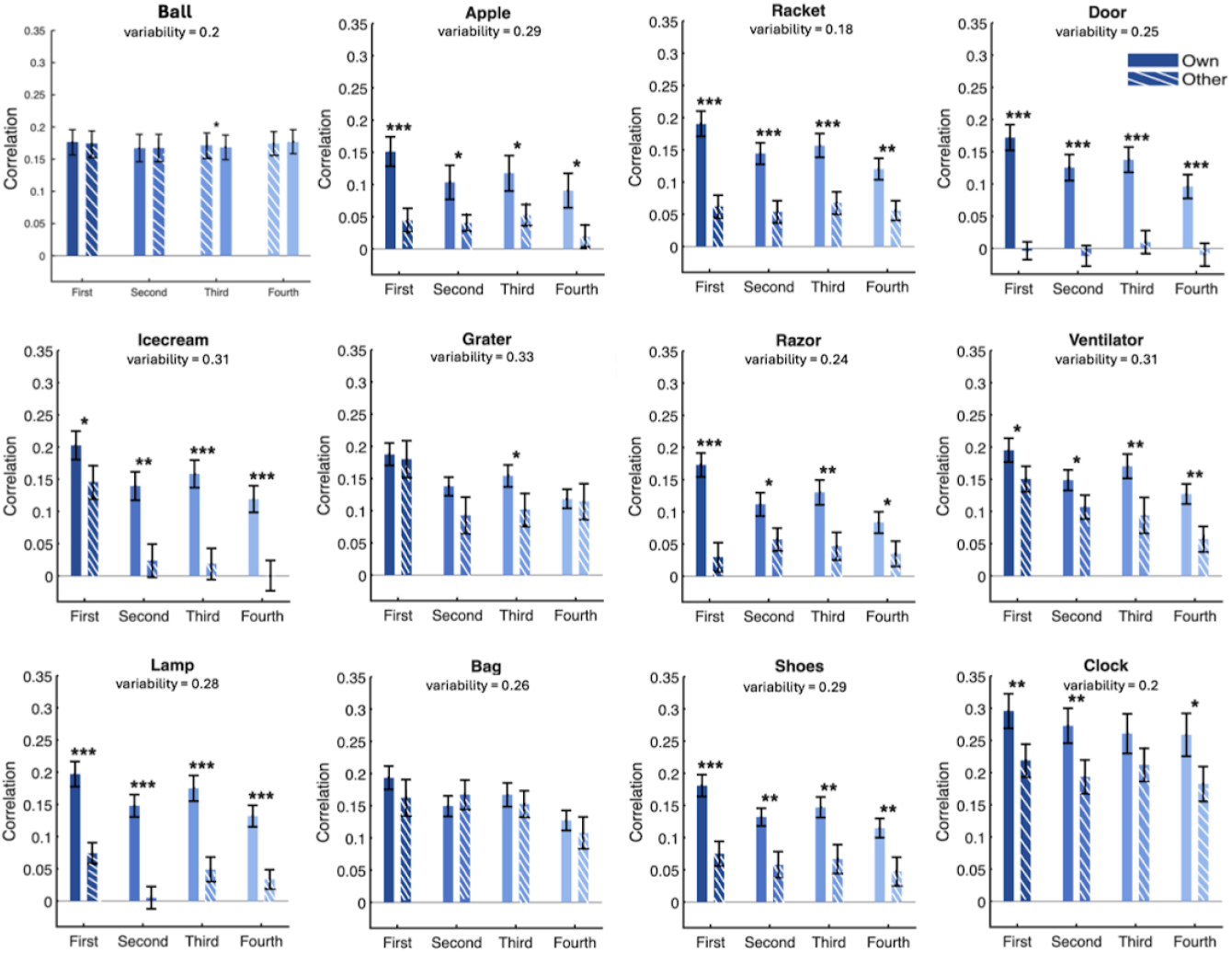
Correlation between individual target similarity and RT for each object category in Experiment 2. For each category, we computed the cosine similarity between the VGG16 activation vectors for each participant’s own drawing of an object and activation vectors for all target images of the corresponding category in the search task. We then correlated these similarity values and the average RT across participants for that object in the corresponding target-present trials (within-participant correlation, “own” condition) and compared it to the same correlations when other participants’ object drawings were used (between-participant correlation, “other” condition). Variability indicates the average pairwise dissimilarity of object exemplar drawings across participants (see Fig. 2A). Error bars indicate SEM. *: p<0.05, **: p<0.01, ***: p<0.001(FDR-corrected where applicable).

